# Genomic evolution of epitopes and Low Complexity Regions in Plasmodium

**DOI:** 10.1101/2021.01.29.428855

**Authors:** Sarah N. Medley, Alyssa Beaudet, Helen Piontkivska, Fabia U. Battistuzzi

## Abstract

Despite decades-long efforts to eradicate malaria, pathogens genomic complexity and variability continue to pose major challenges for the vaccine and drug development. Here we examined the evolutionary history of epitopes and epitope-like regions to determine whether they share underlying evolutionary mechanisms and potential functions that are relevant to pathogens interactions with the host immune response. Our comparative sequence analyses contrasted patterns of sequence conservation, amino acid composition, and protein structure of epitopes and low complexity regions (LCRs) in 21 Plasmodium species. Our results revealed many similarities in amino acid composition and preferred secondary structures between epitopes and LCRs; however, we also identified differences in evolutionary trends where LCRs exhibit overall lower sequence conservation and higher disorder. We also found that both epitopes and LCRs have a wide array of configurations, with various levels of sequence conservation and structural order. We propose that such combination of different levels of conservation and structural order between epitopes and LCRs in the same gene play a role in maintaining the functional integrity required by the pathogen along with the variability necessary to evade the host immune response, with LCRs playing a role in the evasion particularly in the vicinity of conserved epitopes. Overall, our results suggest that there are at least two categories of LCRs, where some LCRs play a potential protective role for conserved (ordered) epitopes because of their variable (or disordered) sequence, while others are less disordered and are as conserved as epitopes. The former ones may be an evolutionary necessity for Plasmodium to maintain the diversity of epitopes, while the latter category may serve currently unknown function(s) and deserve to be examined in greater detail. Our findings show that there may be many more candidate targets for future anti-malarial treatments than initially thought and that some of these targets may work across strains and species.

## Introduction

The decades-long research efforts to eradicate malaria have produced a number of drugs effective against the *Plasmodium* pathogens and many vaccine candidates. Of these vaccines, the only one currently being used in a pilot study is RTS,S but its effectiveness is sub-optimal (Neafsey et al. 2015; Stanisic et al. 2013; RTS,S Clinical Trials Partnership 2015; Olotu et al. 2016). The challenge of vaccine development is the identification of a target that is stable, easily accessible, and that elicits an immunogenic response in the host after its recognition by the immune system. Therefore, comparative sequence and structural analyses of a pathogen’s genome can provide key information on the role of known and new sequences as vaccine targets (Mitran & Yanow 2020).

From a molecular perspective, targets of current vaccines are epitopes, which are short, often repetitive subsequences of an antigen that are exposed to the surface of cells once translated. However, variability in epitope sequence and structure hinder our ability to create effective and long-lasting immunological responses to vaccines (Hou et al. 2020; Anderson et al. 1994). In *Plasmodium*, research efforts have produced many valuable insights on its epidemiology and genome evolution but the evolutionary history of epitopes, and similar subsequences, is still largely unknown. To improve the development of antimalarial drugs it is therefore important to explore the evolutionary history of epitopes and similar sequences that could improve the strength and reliability of current targets.

Here we explore the evolutionary history of epitopes and epitope-like regions to determine possible shared evolutionary mechanisms and candidate functions. In particular, we analyze trends in low complexity regions (LCRs) that are common features of eukaryotic genomes with an especially high frequency in *Plasmodium* (Haerty & Golding 2010; Pizzi & Frontali 2001; Battistuzzi et al. 2016; DePristo et al. 2006; Zilversmit et al. 2010). In most cases, these regions have no known function, although a few hypotheses have been suggested (e.g., tRNA sponges, immune evasion) (Hou et al. 2020; Frugier et al. 2010; Kebede et al. 2019). However, many studies have identified diverse functions for specific LCRs in other organisms, such as altering transcriptional rates or modulating protein-protein interactions (Coletta et al. 2010; Strazic Geljic et al. 2020; María Velasco et al. 2013; Chong et al. 2018). Interestingly, comparisons of LCRs and epitope sequences show that they share some similarities (e.g., amino acid composition, repetitiveness, disorder), which raises some new questions regarding the function and interactions of different genomic regions: (i) is there a correlation between sequence conservation and immunogenic functions? (ii) Are LCR sequences substantially different from other sequences known to be functionally active? (iii) What is the distribution of evolutionary rates in LCRs and what could be their relation to function? (iv) Are the functions of epitopes dependent on the presence of other genomic regions? The first question can be addressed by comparing across species the evolutionary history of epitopes with known immunogenic roles to determine if there is a consistent pattern of conservation. Answers to the second question can be obtained from a direct comparison of sequence characteristics (e.g., composition, structure) between LCRs and epitopes. Insight into the third question can come from an evolutionary analysis of LCRs across *Plasmodium* species in order to identify the level of sequence conservation among these regions and guide possible functional interpretations. Finally, comparisons of associations of epitopes and LCRs can highlight possible functional connections. In particular, we explore two scenarios: LCRs that are highly conserved and LCRs that are variable. Based on the evolutionary arms race between pathogen and host, we argue that both these sets of regions may hold an important piece to the malaria puzzle.

Thus, we have conducted comparative sequence analyses of epitopes and LCRs in 21 *Plasmodium* species (30 strains). Specifically, we focused on sequence conservation, amino acid composition, and protein structure of 2,124 epitopes and 1,062 LCRs to determine shared and unique evolutionary patterns. We identified similarities in sequence and structure between epitopes and LCRs such as amino acid composition and preferred protein domains, but also found differing evolutionary trends with overall lower sequence conservation in LCRs. We found that epitopes (and LCRs) have a wide array of configurations, with various levels of sequence conservation and structural order. We propose that the combination of different levels of conservation and structural order between epitopes and LCRs in the same gene may be a potential mechanism to maintain the functional integrity required by the pathogen along with the variability necessary to evade the host immune response. At the same time, the consistent presence of conserved epitopes and LCRs suggests that there could be potentially new (or expanded) candidate target regions for future treatments.

## Methods

### Epitope identification

From the Immune Epitope Database (IEDB) we collected the complete list of linear epitopes with positive T cell and B cell assay results for epitopes present in at least 2 Plasmodium species or strains out of those represented in IEDB (21 species: *P. vivax, P. inui, P. cynomolgi, P. fragile, P. knowlesi, P. coatneyi, P. ovale, P. malariae, P. berghei, P. yoelii, P. vinckei, P. chabaudi, P. adleri, P. gaboni, P. blacklocki, P. billcollinsi, P. reichenowi, P. praefalciparum, P. falciparum, P. gallinaceum, and P. relictum;* 30 strains: *P. vivax 01, P. vivax Sal 01, P. inui San Antonio 1, P. cynomolgi B, P. cynomolgi M, P. fragile nilgiri, P. knowlesi H, P. knowlesi Malayan PK1 A, P. coatneyi Hackeri, P. ovale curtisi GH01, P. malariae UG01, P. berghei ANKA, P. yoelii yoelii 17XNL, P. yoelii yoelii 17X, P. yoelii yoelii YM, P. vinckei vinckei strain vinckei, P. vinckei vinckei petteri CR, P. chabaudi chabaudi, P. adleri G01, P. gaboni SY75, P. gaboni G01, P. blacklocki G01, P. billcollinsi G01, P. reichenowi CDC, P. reichenowi G01, P. praefalciparum G01, P. falciparum 3D7, P. falciparum IT, P. gallinaceum 8A, and P. relictum SGS1-like*). This search resulted in 569 epitopes present in 7 Plasmodium proteins (AMA1, CP, Enolase, GAPDH, LDH, MSP1, SSP2). We merged epitopes of the same assay type in the same species that overlapped by 50% or more into one to obtain a new total of 167 epitope regions. Because not every Plasmodium species is equally represented in IEDB, we used conservation levels in multiple species alignments to computationally identify epitopes in other species. After aligning each gene across the 21 species (see below), we identified regions in other species in the alignment with at least 50% identity to the epitope regions from IEDB and added them to our list of epitopes. This produced a new list of computationally-identified epitopes (2,124) based on those with confirmed positive assay.

### LCR identification

We identified LCRs with SEG using a window size of 15 but with varying complexity thresholds from 0 to 2.5. All 7 proteins that contained epitopes in multiple species also contained LCRs in multiple species. We found 1,062 total LCRs (1 with complexity 0, 1 with complexity 0.5, 14 with complexity 1, 76 with complexity 1.5, 203 with complexity 2, and 767 with complexity 2.5).

### Conservation

Homologous gene sequences from PlasmoDB were collected for each of our genes and gene alignments were created using MEGA X (v 10.0.4) (Kumar et al. 2018). These alignments were used to calculate the Jensen-Shannon divergence (JSD) method to generate a conservation score at each site (Capra & Singh 2007). Any sites that could not have a score calculated because of too many gaps in the alignment (≥ 30%) were ignored in all of the following conservation calculations. For every gene, we obtained a protein background conservation score by averaging the scores from each site and calculating its standard deviation (stdev). We separately calculated a conservation score for each epitope and LCR by averaging across the scores at each site in the epitope or LCR region. We then used the background score ± 1 stdev as thresholds for epitope and LCR regions to be considered nonconserved (if lower than the background score - 1 stdev) and conserved (if higher than the background score + 1 stdev). Any epitope/LCR scores that fell in between this range was considered to be at the background level of the protein. We also recalculated the JSD scores for individual clades and for species sharing the same host category (human, non-human primate, rodent, bird/reptile). In the analysis of conservation by site, only genes with at least 30 epitope and 30 LCR sites were considered (AMA1, CP, MSP1, and SSP2). The percentages of conserved sites in epitopes and LCRs were compared using a two-proportions Z-test (two-sided).

### Amino acid composition

We determined the percent frequency of amino acids for each of the 30 species/strains within the epitope and LCR regions and for the entire proteome. We compared the average percent frequency of each amino acid for epitopes vs. the proteome and epitopes vs. LCRs. In order to compare the averages, Shapiro-Wilk tests and Q-Q plots were used to assess whether the distribution of percent frequencies were normally distributed across species. The average percent frequencies were then compared using paired t-tests or Wilcoxon signed rank tests (two-sided) depending on whether there was significant evidence for nonnormality. The total percent frequency of amino acids based on their type (nonpolar/aliphatic: Ala, Gly, Ile, Leu, Met, and Val; polar/uncharged: Cys, Asn, Pro, Gln, Ser, and Thr; aromatic: Phe, Trp, and Tyr; positively charged: His, Lys, and Arg; negatively charged: Asp and Glu) and nucleotide composition (AT rich: Phe, Ile, Lys, Leu, Met, Asn, and Tyr; GC rich: Ala,Gly, Pro, Arg, and Trp; balanced: Cys, Asp, Glu, His, Gln, Ser, Thr, and Val) were also determined by species for epitopes, LCRs, and the proteome.

### Protein domains

We used Interpro to obtain protein domain data for each of the 7 proteins in each species independently (Blum et al. 2021). Sites were only considered in further analysis if they had a domain type that agreed in at least 50% of the species in the alignment. In addition, genes that did not have at least 30 epitope and 30 LCR sites were excluded from the analysis. An epitope or LCR region was considered to be disordered if it contained at least 3 disordered sites.

### Overlapping and hyperconserved LCRs

Using the measures described above, we independently analyzed also the regions of LCRs and epitopes that are overlapping and compared these patterns to those in epitopes or LCRs separately. Only the sites shared by the epitope and LCR were included in this analysis. We also analyzed separately a subset of 266 LCRs that showed an unusually high level of sequence conservation. These were identified as the top 25% LCRs with JSD conservation score higher than the background conservation score+1 stdev. The percentages of extracellular membrane bound sites in epitopes and hyperconserved LCRs were compared using a two-proportions Z-test (two-sided).

## Results

The efficacy of antimalarial vaccines is dependent, in part, on the quality and accessibility of the target sequence. Desirable properties in a target include (i) conservation, which supports long-term stability, (ii) accessibility, for ease of targeting, (iii) specificity, which allows sequences to be targeted uniquely, and (iv) the presence of adjuvant targets that could strengthen and refine the targeting process (MacRaild et al. 2018). For the epitopes of *Plasmodium* species, (ii) and (iii) are relatively well known but (i) and (iv) are not. These properties that make a sequence a good vaccine target are the opposite of those that would protect the pathogen from detection by the immune system. This inverted relation raises an interesting evolutionary question: how variable can target sequences in *Plasmodium* be and what does it mean for the development of future treatments/prevention strategies? To answer these questions, we performed a series of comparative genomic analyses of epitopes shared by multiple *Plasmodium* species to assess their properties as targets and computationally identify potential adjuvant or new targets. The comparative genomic approach we use allows us to reconstruct the evolutionary process of these regions in multiple genomes, irrespective of the fact that they may currently act or be identified as epitopes.

### Dataset

From 21 species and 30 strains of *Plasmodium*, which included primate, rodent, and bird pathogens, we identified 2,124 epitopes shared by at least 2 genomes (either 2 species or 2 strains). These epitopes were located in 7 genes (Table 1) that are known for their functions in parasite blood (AMA1, MSP1) and liver stages (CP, SSP2) as well as metabolism (Enolase, GAPDH, LDH). We found that all 7 genes also contained 1,062 LCRs in at least 2 genomes.

**Table 1.** Frequencies of conservation of epitope and LCR sites per Plasmodium clade. Percentage show the average of multiple species within each clade. Clade size: Plasmodium 11, Vinckeia 7, Laverania 10, Haemamoeba 2.

|  | % Epitope<br>Conserved<br>Sites | % Epitope<br>Background<br>Sites | % Epitope<br>Nonconserved<br>Sites | % LCR<br>Conserved<br>Sites | % LCR<br>Background<br>Sites | % LCR<br>Nonconserved<br>Sites |
| --- | --- | --- | --- | --- | --- | --- |
| Plasmodium | 25.1 | 64.0 | 10.9 | 12.7 | 63.4 | 23.9 |
| Vinckei | 34.4 | 59.4 | 6.3 | 9.5 | 60.4 | 30.2 |
| Laverania | 23.7 | 63.0 | 13.3 | 9.9 | 64.8 | 25.3 |
| Haemamoeba | 37.7 | 58.1 | 4.1 | 11.6 | 64.4 | 24.1 |

### Epitope sequence and structure composition

Compositional trends in epitopes relative to the proteome can provide information on the evolutionary forces these regions are exposed to. We focused on the 8 amino acids that are most commonly used in epitopes (> 5% frequency; **Table S1**) and compared their usage in epitopes *vs.* the genes they are embedded in and *vs.* the overall proteome. We found that all of these amino acids are also the most commonly used in the proteome, with the exception of Gly (and all have >5% frequencies in the genes). However, the relative percent use of each amino acid differs with 6 of these amino acids having significantly different frequencies when compared to the individual genes (Asp, Glu, Ile, Lys, Leu, Asn) and also to the proteome (Asp, Glu, Gly, Ile, Asn, Ser). The difference between genes and proteomes relative to epitopes could be explained by the usage of Ser, Gly, and Lys in LCRs (see below). While these results would seem to suggest that the evolutionary pressures acting on epitopes are optimized for their functions, we also found a large variance in amino acid frequencies among species and among epitopes, which weakens our ability to identify generalized trends.

In agreement with previous studies that have found that epitopes preferentially use hydrophilic amino acids, we found that of the 8 most commonly used amino acids, which account for ~58% of the epitope composition, 5 are hydrophilic (Asn, Lys, Asp, Glu, Ser) and three are hydrophobic (Leu, Ile, Gly) (**Figure 1A, Table S1**) (Soga et al. 2010). Both hydrophilic and hydrophobic amino acids show statistically significantly different frequencies between epitopes and genes/proteome. While the reason for these differences is unclear, they show that both categories of amino acids are equally important in the evolutionary history of epitopes (Pan et al. 2018; Chowell et al. 2015). Finally, we note that patterns in amino acid composition do not show any significant difference among the four major clades of Plasmodium (Plasmodium, Vinckeia, Laverania, Haemamoeba).

**Figure 1.**
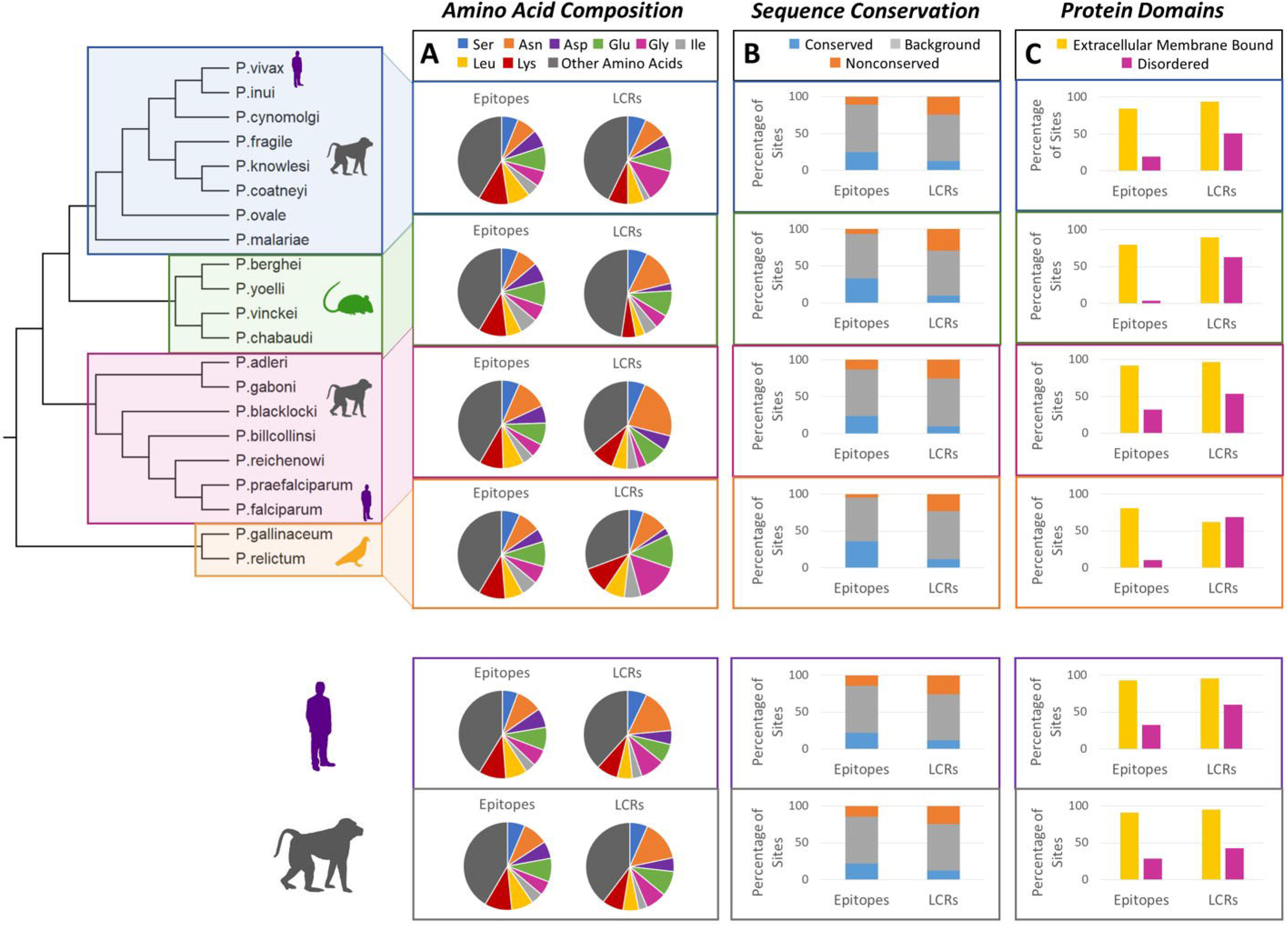
Epitope and LCR sequence and structural features across phylogenetic subgroups and host preference. A) amino acid frequency (%). B) sequence conservation (JSD). C) predicated protein domains (interpro).

In terms of structure, epitopes have ≥ 90% extracellular membrane bound sites and ~30% sites that are in a disordered configuration. The cell surface protein (CP, SSP2) epitopes have especially high frequencies of disorder (66% in CP and 56% in SSP2) which is interesting considering that, for CP, these are the sites targeted by RTS,S. In contrast, the epitopes in the cell membrane proteins (AMA1, MSP1), as well as the metabolism gene GAPDH, have no identified disordered sites (**Figure S1**).

### Epitope sequence and structure conservation

Using our definition of conserved, background, and non-conserved regions (see Methods) we found ~20% of epitopes to be conserved, 75% to be background, and 5% to be non-conserved (**Figure 1B**). A similar result was obtained for all sites individually, instead of averaging over the epitope regions. Among genes, we observe comparable levels of conservation (17-25% conserved epitope sites) although the pathogenicity-related genes (AMA1, CP, MSP1, and SSP2) have a higher frequency of non-conserved epitope sites (~20%; **Figure 2A**). A similar pattern is found among clades. The exceptions are Vinckeia and Haemamoeba that show higher frequencies of conserved epitopes and lower frequencies of non-conserved (Table 1). It is unclear if this trend can be attributed to an evolutionary or biological uniqueness of these species or to a small sample size artifact. The observed distribution of background and conserved epitopes is unexpected, especially in genes that should favor antigenic diversity to prevent detection from the immune system of the host and that, therefore, should have a higher percentage of non-conserved (variable) epitopes (Pilosof et al. 2019). Even from a structural point of view, epitopes do not necessarily follow expectations. While the vast majority of them in all species and genes are composed of extracellular sites, the exposure to an outside solvent would predict a high percentage of disorder (MacRaild et al. 2018). Instead, we find that, on average, ~ 30% of the epitope sites are disordered and in Haemamoeba and Vinckeia this percentage is even lower (11% in Haemamoeba and 4% in Vinckeia; **Figure 1C**).

**Figure 2.**
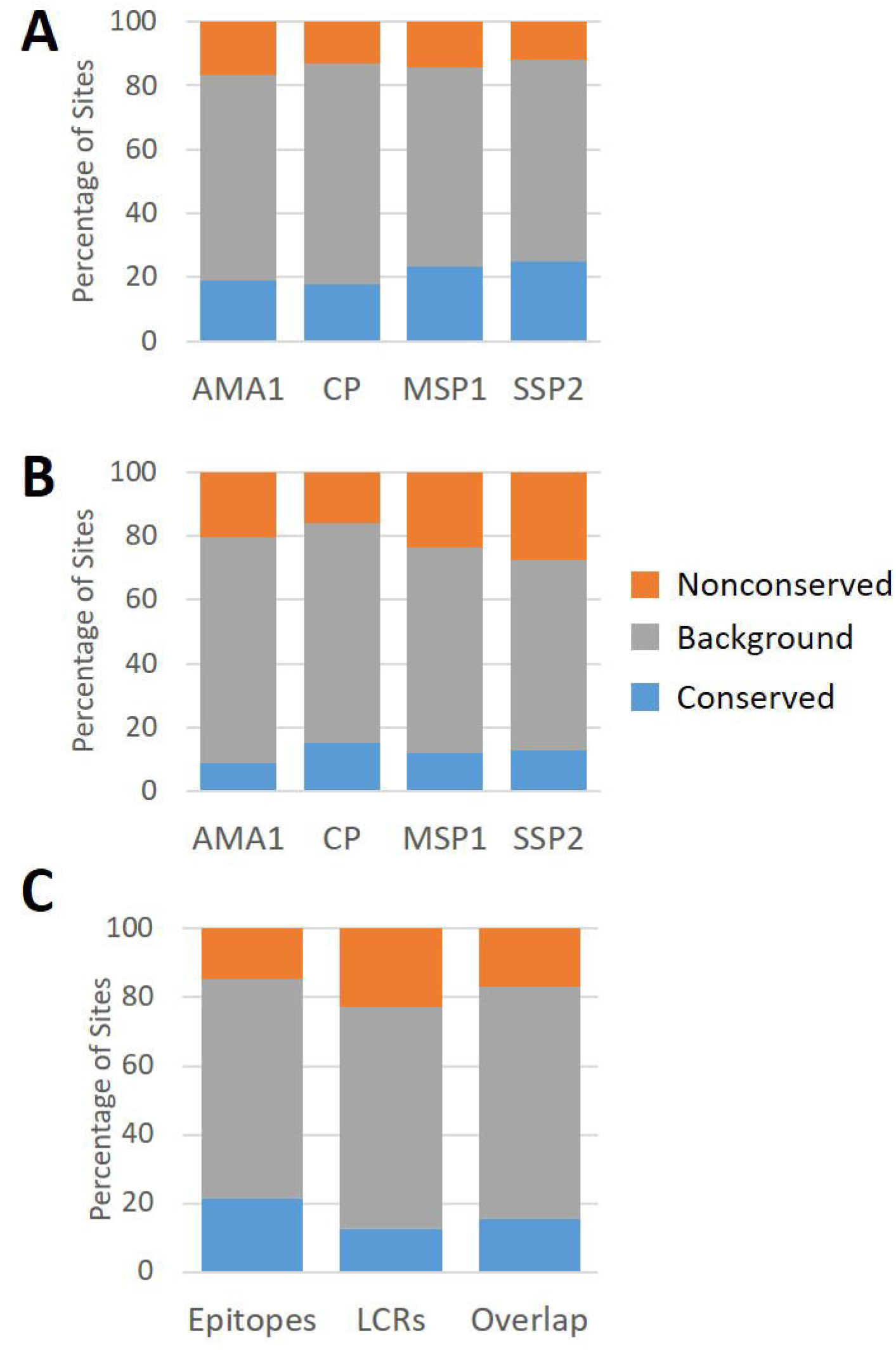
Proportion of conserved, background, and nonconserved sites by gene (A: epitopes and B:LCRs) and for different types of regions (C).

### Low complexity sequences as epitope-like regions

We found that 33% of epitopes overlap with an LCR by at least one amino acid resulting in a total of 778 epitope-LCR pairs with a median overlap of 13 amino acids. Thus, we expanded our investigations to LCRs (those that are overlapping and those that are not) and compared these results to those found in epitopes to assess the extent of similarity between these two regions. The strongest difference is the level of conservation. In a site-by-site analysis, we found LCRs and epitopes to have the majority (~60%) of their sites with the same background sequence conservation as the rest of the protein (**Figure 2C**, grey bars), but LCRs have a lower frequency of conserved sites compared to epitopes (proportion test p-value ≪ 0.001, two-sided) (**Figure 2B and C**). Out of all LCRs, we found 11% to be conserved compared to 20% of epitopes (Table S3). This result is not surprising since many previous studies have shown that LCRs are fast evolving regions with high mutation rates (Huntley & Clark 2007; Haerty & Golding 2011; Battistuzzi et al. 2016; Chaudhry et al. 2018; Lenz et al. 2014; Wang & Harrison 2020). However, LCR sites overlapping with epitopes show higher conservation than the nonoverlapping sites suggesting that the evolution of these regions is driven by the epitopes (proportion test p-value = 0.02, two-sided) (**Figure 2C**).

Despite differences in conservation, of the 8 amino acids that account for 75% of the total composition, LCRs share five highly used amino acids with epitopes (Asn, Glu, Gly, Lys, Ser) while another three are preferentially used only in LCRs (Ala, Pro, Thr). The preference for these last three amino acids is not surprising considering all three are found in high frequency in disordered proteins (see below; Theillet et al. 2013; Coskuner-Weber & Uversky 2019; Perez et al. 2014).

Interestingly, the relative use of these amino acids among species is more variable in LCRs compared to epitopes with 4 of the five amino acids that are shared showing significantly different average frequencies (Wilcox p-value ≪ 0.05 for Lys, Asp, Ile, Leu). Even those amino acids that overall do not appear to be used at different frequencies in LCRs and epitopes, like Asn, show significant differences when analyzed by clade (in particular the Laverania clade has a significantly higher usage of Asn in LCRs, Wilcoxon p-value ≪ 0.001, respectively) (**Figure 1A**). This is in agreement with previous studies that have shown the prevalence of Asn in these clades (e.g., Zilversmit et al. 2010; Chaudhry et al. 2018). In terms of protein domains, LCRs and epitopes share a very high percentage of extracellular membrane bound sites. LCRs have a higher overall proportion of disordered sites (~50%) compared to epitopes, even though this percentage is lower in cell membrane proteins (AMA1: 0% and MSP1: 25%), like we have observed in epitopes (and is higher in CP and SSP2 like in epitopes).

Interestingly, despite the general low sequence conservation of LCRs, we identified a subset of them with conservation values greater than the background gene level and similar to epitope conservation. We refer to this subset of LCRs as hyperconserved LCRs. Approximately 10% of epitopes overlap with a hyperconserved LCR and, accordingly, these epitopes show an overall higher sequence conservation than others. These two regions also share similar amino acid composition with the hyperconserved LCRs using the same 8 amino acids consistently across species (like epitopes). However, the hyperconserved LCRs are different from epitopes in terms of structure showing approximately half of the amount of disorder (~15%) of epitopes (and ¼ of that in other LCRs) all in a single gene (SSP2) (**Figure S1**). They also have slightly but significantly fewer extracellular membrane bound sites compared to epitopes (proportion test p-value = ≪0.001, two-sided). Even considering this difference, the majority (~80%) of the sites are still extracellular, making these highly conserved regions potentially accessible and stable targets. The differences between hyperconserved and other LCRs and the similarities of the former ones to epitopes suggest that the hyperconserved LCRs may be a subcategory of LCRs with potentially different functions.

### Effect of phylogenetic ancestry and host preference on epitope and LCR evolution

In order to determine if any differences between epitopes and LCRs could be explained by ancestry or selective pressures from different host immune systems, we compared the characteristics of epitopes and LCRs across clades (phylogenetic ancestry) and by host preference of the pathogen (**Figure 1 upper and lower panels**). Although sequence conservation remains low, sequence composition and structural features are conserved for epitopes across all clades and between all host preferences. The only major difference is the low proportion of disordered sites in the Vinckeia and Haemamoeba clade. LCRs instead have similar levels of disorder in all groups irrespective of their ancestry or host preference, but Haemamoeba again show unique trends in the number extracellular membrane sites. It is possible that the different trends observed in Vinckeia and Haemamoeba are driven by the small number of species within these two groups (3 and 2 respectively). Additionally, LCRs change drastically in their amino acid composition between clades, particularly in their usage of the amino acids Asn and Gly (see above). Overall, there are more differences between clades than by host preference, suggesting that evolutionary history rather than host choice has a stronger effect on epitope and LCR evolution.

## Discussion

Our study analyzed the extent of variation in epitopes and compared their properties to those of other common regions in Plasmodium with the goal of understanding the role of sequence and structural conservation in relation to immunogenicity. We found that Plasmodium genomes have various types of epitopes with composition, structure, and conservation properties that are often similar to those of LCRs. The variability in the epitope (and LCR) properties is difficult to explain with a single evolutionary mechanism and we propose that interactions between epitopes and LCRs may provide a new perspective on it. Starting from sequence conservation, epitopes that are non-conserved have been found to be favored by within-host competition of multiple strains and multiple species of Plasmodium (Richie 1988; Pilosof et al. 2019). Accordingly, we found that the majority of the epitopes we analyzed are non-conserved or follow the same conservation level of their proteins. However, we also found that ~20% of epitopes are conserved. These regions are potentially problematic for the pathogen because they are more easily recognized by the host immune system, especially during co-infections (Pilosof et al. 2019). Thus, we would expect that the pathogen would have evolved a mechanism to mask these conserved regions. We propose that this mechanism may involve LCRs.

LCRs are known for their often disordered form and high evolutionary rates (Haerty & Golding 2011; Zilversmit et al. 2013, 2010; Chaudhry et al. 2018). The LCRs that we identified in the same genes as epitopes follow these properties, being less conserved than epitopes (11% vs. 20%) and more disordered (50% vs. 30%). Interestingly, the LCRs that are located farther away from conserved epitopes are five times more likely to be disordered than those that are either overlapping or close to an epitope (8% vs. 1.5%), while variable epitopes are more likely to correlate to non-disordered distant LCRs compared to overlapping ones (70% vs. 32%) (**Table 2**). Similarly, there is a higher correlation between variable LCRs and ordered epitopes for distant pairs than for close pairs (85.4% vs. 30.4%) (**Table S2**). We hypothesize that the pairing of conservation with disorder (and order with non-conservation) may facilitate a conformational masking strategy of LCRs on epitopes that would otherwise be readily recognized by the immune system (Kwong et al. 2002; Goo et al. 2012; Morales et al. 2015; Davies et al. 2017; Herrera et al. 2015). Variable (or disordered) epitopes, instead, do not require the “protection” of LCRs because they can evade the immune system through sequence variation. This model would explain how the pathogen can maintain in its genome both conserved/non-conserved and ordered/disordered epitopes.

**Table 2.** Correlation of conservation and disorder in epitope-LCR pairs. Similar trends are seen in near (within 10 amino acids) and distant (> 10 amino acids) pairs (Table S2).

|  |  | Non-disordered<br>LCRs | Disordered<br>LCRs |
| --- | --- | --- | --- |
| Overlapping pairs | Variable epitopes | 245 (31.5%) | 359 (46.1%) |
|  | Conserved epitopes | 162 (20.8%) | 12 (1.5%) |
|  |  | Non-disordered<br>LCRs | Disordered<br>LCRs |
| Non-overlapping pairs | Variable epitopes | 1101 (70.4%) | 115 (7.3%) |
|  | Conserved epitopes | 224 (14.3%) | 125 (8.0%) |

However, even with this model, another question remains: why do some epitopes form ordered structures and others do not? To answer this question, we need to consider two variables: amino acid composition and the role of epitopes in immune evasion. For amino acid composition, previous studies(Soga et al. 2010) have suggested that epitopes are preferentially composed of hydrophilic amino acid, which are known to promote disorder in proteins (Weathers et al. 2004). However, we found that epitopes are composed preferentially of 8 amino acids, three hydrophobic (Leu, Ile, Gly) and five hydrophilic (Asn, Lys, Asp, Glu, Ser) that are primarily within ordered structures (~70%). Thus, the composition of epitopes cannot fully explain their structural properties.

We propose that the explanation for having both ordered and disordered epitopes in Plasmodium may be related to host immune response. Disorder in LCRs has been often explained as a strategy that promotes immune evasion and that, at the same time, makes them bad targets for vaccines because of lack of immune response (Mendes et al. 2013). The fact that we found a number of disordered epitopes known to trigger immunogenic responses suggests that this correlation between disorder and lack of immune response is too simplistic. A recent study by MacRaild et al (2018) offers a new interpretation. In this study the authors have found a positive correlation between disorder in epitopes and positive antibody binding assay suggesting that the absence of a secondary structure does not prevent recognition from the antibody and, thus, an immune response. If confirmed, this would allow the pathogen to use either ordered or disordered epitopes based on its needs for immune evasion and using LCRs as a protective variable sequence (**Table S2**). Moreover, because many LCRs are also disordered, it is possible to speculate that some LCRs, especially those that are overlapping with epitopes, may be used as adjuvant targets. This is an intriguing option since we found that 33% of epitopes overlap with an LCR and 45% of epitopes have an LCR within 10 amino acids. If these LCRs were found to be immunogenic, it would significantly expand the size and numbers of candidate target sequences.

Overall, our results suggest that there are at least two categories of LCRs, those that because of their variable (or disordered) sequence may play a protective role for conserved (ordered) epitopes and those that are as conserved as epitopes (and less disordered). The former ones may be an evolutionary necessity for Plasmodium to maintain the diversity of epitopes it has (although the ultimate reason for this diversity is still unclear). The latter ones suggest that these LCRs may be worth analyzing as functional units. In either of these cases, though, our results show that there may be many more candidate targets for future anti-malarial treatments than initially thought and that some of these targets may work across strains and species.

## Acknowledgments

This work was supported by funds from Oakland University and from the National Institute of Health [grant number R15GM121981] to FUB.

## References

Anderson DE, Malley A, Benjamini E, Gardner MB, Torres JV. 1994. Hypervariable epitope constructs as a means of accounting for epitope variability. Vaccine. 12:736–740. doi: 10.1016/0264-410x(94)90225-9.

Battistuzzi FU et al. 2016. Profiles of low complexity regions in Apicomplexa. BMC Evolutionary Biology. 16:47.

Blum M et al. 2021. The InterPro protein families and domains database: 20 years on. Nucleic Acids Research. 49:D344–D354. doi: 10.1093/nar/gkaa977.

Capra JA, Singh M. 2007. Predicting functionally important residues from sequence conservation. Bioinformatics. 23:1875–1882. doi: 10.1093/bioinformatics/btm270.

Chaudhry SR, Lwin N, Phelan D, Escalante AA, Battistuzzi FU. 2018. Comparative analysis of low complexity regions in Plasmodia. Sci Rep. 8:1–9. doi: 10.1038/s41598-017-18695-y.

Chong S et al. 2018. Imaging dynamic and selective low-complexity domain interactions that control gene transcription. Science. 361. doi: 10.1126/science.aar2555.

Chowell D et al. 2015. TCR contact residue hydrophobicity is a hallmark of immunogenic CD8+ T cell epitopes. PNAS. 112:E1754–E1762. doi: 10.1073/pnas.1500973112.

Coletta A et al. 2010. Low-complexity regions within protein sequences have position-dependent roles. BMC Systems Biology. 4:43. doi: 10.1186/1752-0509-4-43.

Coskuner-Weber O, Uversky VN. 2019. Alanine Scanning Effects on the Biochemical and Biophysical Properties of Intrinsically Disordered Proteins: A Case Study of the Histidine to Alanine Mutations in Amyloid-β42. J. Chem. Inf. Model. 59:871–884. doi: 10.1021/acs.jcim.8b00926.

Davies HM, Nofal SD, McLaughlin EJ, Osborne AR. 2017. Repetitive sequences in malaria parasite proteins. FEMS Microbiology Reviews. 41:923–940. doi: 10.1093/femsre/fux046.

DePristo MA, Zilversmit MM, Hartl DL. 2006. On the abundance, amino acid composition, and evolutionary dynamics of low-complexity regions in proteins. Gene. 378:19–30. doi: 10.1016/j.gene.2006.03.023.

Frugier M et al. 2010. Low Complexity Regions behave as tRNA sponges to help co-translational folding of plasmodial proteins. FEBS Letters. 584:448–454. doi: 10.1016/j.febslet.2009.11.004.

Goo L, Milligan C, Simonich CA, Nduati R, Overbaugh J. 2012. Neutralizing Antibody Escape during HIV-1 Mother-to-Child Transmission Involves Conformational Masking of Distal Epitopes in Envelope. Journal of Virology. 86:9566–9582. doi: 10.1128/JVI.00953-12.

Haerty W, Golding GB. 2011. Increased Polymorphism Near Low-Complexity Sequences across the Genomes of Plasmodium falciparum Isolates. Genome Biol Evol. 3:539–550. doi: 10.1093/gbe/evr045.

Haerty W, Golding GB. 2010. Low-complexity sequences and single amino acid repeats: not just “junk” peptide sequences. Genome. 53:753–762. doi: 10.1139/G10-063.

Herrera R et al. 2015. Reversible Conformational Change in the Plasmodium falciparum Circumsporozoite Protein Masks Its Adhesion Domains. Infection and Immunity. 83:3771–3780. doi: 10.1128/IAI.02676-14.

Hou N et al. 2020. Low-Complexity Repetitive Epitopes of Plasmodium falciparum Are Decoys for Humoural Immune Responses. Front. Immunol. 11. doi: 10.3389/fimmu.2020.00610.

Huntley MA, Clark AG. 2007. Evolutionary analysis of amino acid repeats across the genomes of 12 Drosophila species. Molecular biology and evolution. 24:2598–2609.

Kebede AM, Tadesse FG, Feleke AD, Golassa L, Gadisa E. 2019. Effect of low complexity regions within the PvMSP3α block II on the tertiary structure of the protein and implications to immune escape mechanisms. BMC Struct Biol. 19:6. doi: 10.1186/s12900-019-0104-0.

Kumar S, Stecher G, Li M, Knyaz C, Tamura K. 2018. MEGA X: Molecular Evolutionary Genetics Analysis across Computing Platforms. Mol Biol Evol. 35:1547–1549. doi: 10.1093/molbev/msy096.

Kwong PD et al. 2002. HIV-1 evades antibody-mediated neutralization through conformational masking of receptor-binding sites. Nature. 420:678–682. doi: 10.1038/nature01188.

Lenz C, Haerty W, Golding GB. 2014. Increased Substitution Rates Surrounding Low-Complexity Regions within Primate Proteins. Genome Biology and Evolution. 6:655–665. doi: 10.1093/gbe/evu042.

MacRaild CA, Seow J, Das SC, Norton RS. 2018. Disordered epitopes as peptide vaccines. Peptide Science. 110:e24067. doi: https://doi.org/10.1002/pep2.24067.

María Velasco A et al. 2013. Low complexity regions (LCRs) contribute to the hypervariability of the HIV-1 gp120 protein. J. Theor. Biol. 338:80–86. doi: 10.1016/j.jtbi.2013.08.039.

Mendes T a. O et al. 2013. Repeat-Enriched Proteins Are Related to Host Cell Invasion and Immune Evasion in Parasitic Protozoa. Mol Biol Evol. 30:951–963. doi: 10.1093/molbev/mst001.

Mitran CJ, Yanow SK. 2020. The Case for Exploiting Cross-Species Epitopes in Malaria Vaccine Design. Front. Immunol. 11. doi: 10.3389/fimmu.2020.00335.

Morales RAV et al. 2015. Structural basis for epitope masking and strain specificity of a conserved epitope in an intrinsically disordered malaria vaccine candidate. Scientific Reports. 5:10103. doi: 10.1038/srep10103.

Neafsey DE et al. 2015. Genetic Diversity and Protective Efficacy of the RTS,S/AS01 Malaria Vaccine. New England Journal of Medicine. 373:2025–2037. doi: 10.1056/NEJMoa1505819.

Olotu A et al. 2016. Seven-Year Efficacy of RTS,S/AS01 Malaria Vaccine among Young African Children. New England Journal of Medicine. 374:2519–2529. doi: 10.1056/NEJMoa1515257.

Pan R et al. 2018. Increased Epitope Complexity Correlated with Antibody Affinity Maturation and a Novel Binding Mode Revealed by Structures of Rabbit Antibodies against the Third Variable Loop (V3) of HIV-1 gp120. Journal of Virology. 92. doi: 10.1128/JVI.01894-17.

Perez RB, Tischer A, Auton M, Whitten ST. 2014. Alanine and proline content modulate global sensitivity to discrete perturbations in disordered proteins. Proteins: Structure, Function, and Bioinformatics. 82:3373–3384. doi: https://doi.org/10.1002/prot.24692.

Pilosof S et al. 2019. Competition for hosts modulates vast antigenic diversity to generate persistent strain structure in Plasmodium falciparum. PLOS Biology. 17:e3000336. doi: 10.1371/journal.pbio.3000336.

Pizzi E, Frontali C. 2001. Low-Complexity Regions in Plasmodium falciparum Proteins. Genome Res. 11:218–229. doi: 10.1101/gr.152201.

Richie TL. 1988. Interactions between malaria parasites infecting the same vertebrate host. Parasitology. 96:607–639. doi: 10.1017/S0031182000080227.

RTS, S Clinical Trials Partnership. 2015. Efficacy and safety of RTS,S/AS01 malaria vaccine with or without a booster dose in infants and children in Africa: final results of a phase 3, individually randomised, controlled trial. Lancet. 386:31–45. doi: 10.1016/S0140-6736(15)60721-8.

Soga S, Kuroda D, Shirai H, Kobori M, Hirayama N. 2010. Use of amino acid composition to predict epitope residues of individual antibodies. Protein Engineering, Design and Selection. 23:441–448. doi: 10.1093/protein/gzq014.

Stanisic DI, Barry AE, Good MF. 2013. Escaping the immune system: How the malaria parasite makes vaccine development a challenge. Trends Parasitol. 29:612–622. doi: 10.1016/j.pt.2013.10.001.

Strazic Geljic I et al. 2020. Cytomegalovirus protein m154 perturbs the adaptor protein-1 compartment mediating broad-spectrum immune evasion Rath, S, editor. eLife. 9:e50803. doi: 10.7554/eLife.50803.

Theillet F-X et al. 2013. The alphabet of intrinsic disorder. Intrinsically Disordered Proteins. 1:e24360. doi: 10.4161/idp.24360.

Wang Y, Harrison P. 2020. Homopeptide and Homocodon Levels are Coupled to GC/AT Bias Levels, Intrinsic Disorder Propensity and other Factors Across Diverse Fungi. In Review doi: 10.21203/rs.3.rs-118390/v1.

Weathers EA, Paulaitis ME, Woolf TB, Hoh JH. 2004. Reduced amino acid alphabet is sufficient to accurately recognize intrinsically disordered protein. FEBS Letters. 576:348–352. doi: https://doi.org/10.1016/j.febslet.2004.09.036.

Zilversmit MM et al. 2013. Hypervariable antigen genes in malaria have ancient roots. BMC Evolutionary Biology. 13:110. doi: 10.1186/1471-2148-13-110.

Zilversmit MM et al. 2010. Low-Complexity Regions in Plasmodium falciparum: Missing Links in the Evolution of an Extreme Genome. Mol Biol Evol. 27:2198–2209. doi: 10.1093/molbev/msq108.

