## Supplementary figures and images for "Genomic evolution of epitopes and Low Complexity Regions in Plasmodium"

### Supplemental Figure 1

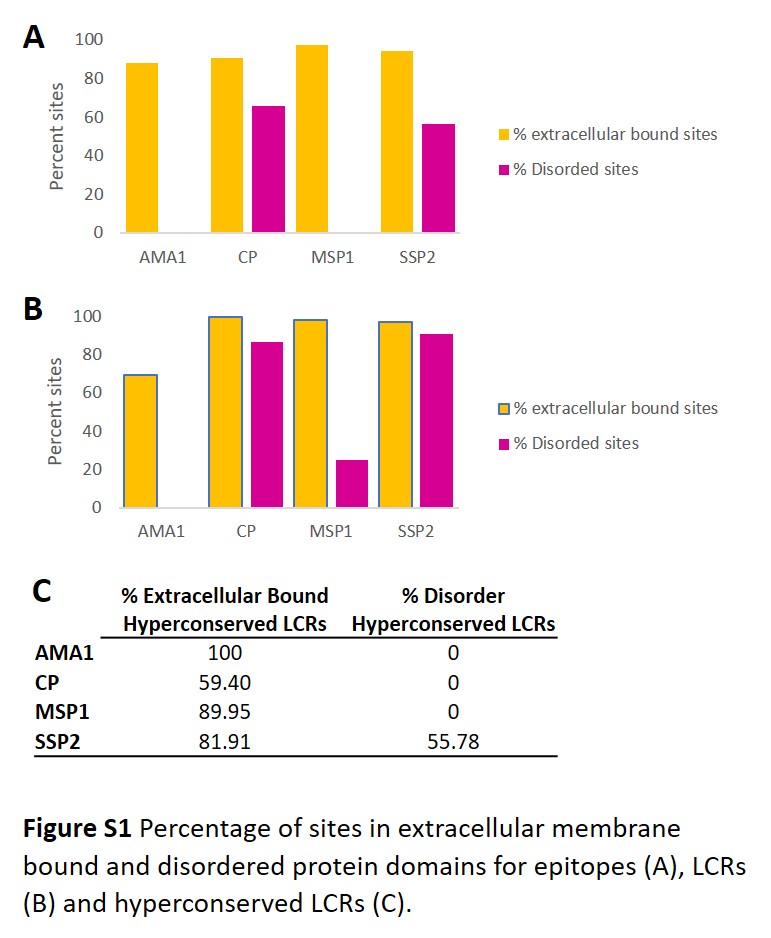

### Supplemental Table 1

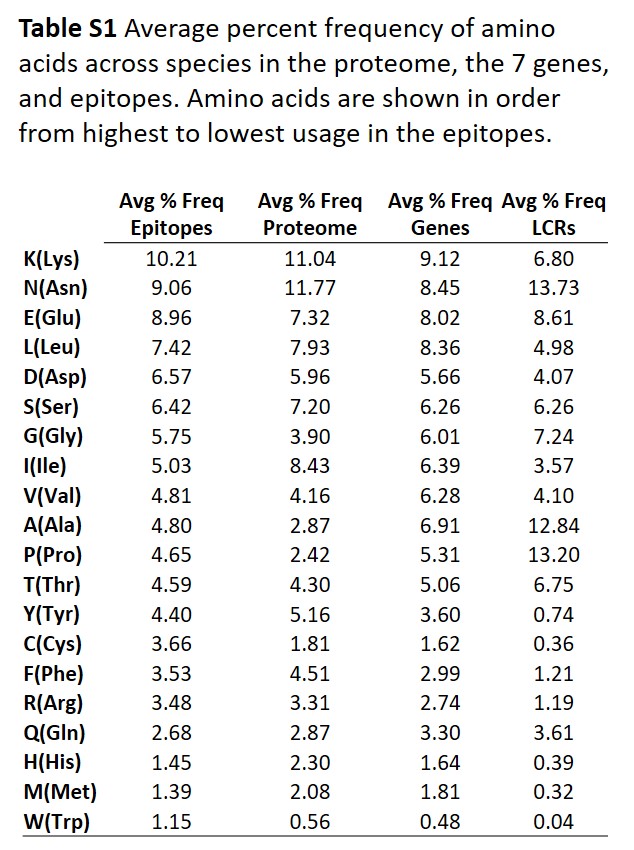

### Supplemental Table 2

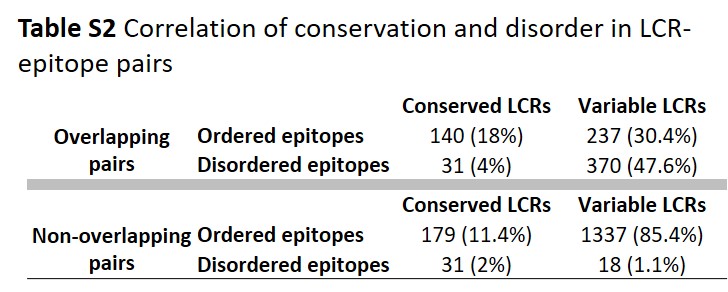
